# Towards Scalable Age-Grading of *Aedes albopictus* mosquito using Mid-Infrared Spectroscopy and Machine Learning

**DOI:** 10.1101/2025.06.20.660660

**Authors:** Mattia Foti, Martina Micocci, Mauro Pazmiño Betancourth, Ivan Casas Gomez-Uribarri, Paola Serini, Beniamino Caputo, Alessandra della Torre, Francesco Baldini

## Abstract

The age structure and dynamics of mosquito populations are crucial for understanding their ability to spread diseases and assessing the effectiveness of anti-mosquito control measures. However, available methods to age-grade mosquito populations are labour-intensive and imprecise, particularly for *Aedes* species. We investigated the potential of Mid-Infrared Spectroscopy (MIRS) combined with Supervised Machine Learning (ML) to rapidly and accurately predict the age of adult females and males of the arbovirus vector, *Aedes albopictus*. First, we demonstrated the ability of MIRS-ML to age male and female mosquitoes reared under laboratory conditions. Second, we optimised the model with adults emerged from wild collected eggs reared under natural conditions in a semi-field facility, to expose them to more realistic ambient conditions. For each sex we developed three ML models based on the resolution of the predicted adult age class: low (9 day interval), medium (6 days) and high resolution (3 days) from 1 to 15 or 33 days for males and females, respectively. The prediction accuracy decreased as the resolution increased. In males, the accuracy dropped from 99% (low) to 93% (medium) and 85.8% (low); in females the high and medium resolution models showed 89.4% and 78.5% accuracy, which decreased to 72.6% for the low resolution. In a simulated vector control intervention, the low-resolution models allowed to detect shifts in the age-structure of *Ae. albopictus* populations with minimal sampling effort (<100 specimens). Finally, we validated MIRS-ML on two unseen data and reconstructed plausible age structures in 1) laboratory-reared and 2) field-collected *Ae. albopictus* males and females. Overall, the results represent a first step towards the development of a sound and reproducible MIRS-ML approach for age-grading of *Ae. albopictus* populations in the wild.

**AUTHOR SUMMARY:** Knowing the age of mosquito populations is critical for understanding how effectively they can transmit viruses like dengue, chikungunya, and Zika, as older mosquitoes are more likely to be infectious. Also, comparing ages of mosquito population before and after a control intervention - such as insecticide aerial spraying - may allow to understand the impact of the intervention. However, existing methods to estimate mosquito age are time-consuming and imprecise. In this study, we tested whether a rapid and scalable method based on detection of age-related changes by mid-infrared spectroscopy (MIRS) coupled with machine learning (ML) could accurately estimate the age of *Aedes albopictus*, the Asian Tiger mosquito, an important arbovirus vector. We trained MIRS-ML models using mosquitoes reared in both laboratory and semi-field conditions to reflect realistic environmental variation. Our models were able to classify mosquito age with high accuracy, especially when grouping individuals into broader age categories. In simulated vector control scenarios, low-resolution models effectively detected shifts in population age structure with minimal sampling effort. We also applied our approach to field-collected mosquitoes that showed plausible age structures, suggesting potential of this approach for real-world surveillance. This method represents a promising, scalable, and non-destructive tool for monitoring mosquito population dynamics and could help monitor control strategies against *Aedes*-borne diseases.

## INTRODUCTION

Mapping the demographic characteristics of wild mosquito populations is a potentially powerful approach to increase accuracy of estimates of the actual risk of mosquito-borne pathogens transmission and to evaluate the output of vector control interventions with a limited sampling effort (1). However, estimates of mosquito age is a difficult task. Traditional methods for estimating age of female mosquito based on changes in ovary morphology related to the gonotrophic cycles) (2,3) are labour-intensive, difficult to standardise, require significant technical expertise and only provide indirect and imprecise estimates of biological age (1,4). In addition to being difficult to apply at a large scale, they have been developed for *Anopheles* mosquitoes, and there is a significant lack of reproducibility when they are applied to other mosquito genera, such as *Aedes* (5,6). Other age-grading methods based on proteomics, gene profiling and gene expression have been proposed in recent years mostly for the major *Anopheles* afrotropical malaria vectors (7,8) and the arbovirus *Aedes* vectors, *Ae. aegypti* and *Ae. albopictus* (4,9–11). However, the large-scale application of these approaches is constrained by lack of conclusive studies on their reproducibility and by the high implementation costs, and none of them has become a gold standard for mosquito population age-grading (12,13).

In recent years, Infrared spectroscopy (IR) has been proposed as an alternative for mosquito age-grading based on the detection of age-related changes in the chemical composition of the mosquito’s cuticle (14). By directing light onto a sample, IR quantifies the energy absorbed by molecules based on their vibrational modes, providing chemical spectra within seconds and without the need for any reagent (15,16). The age-related spectral changes are then disentangled by machine learning (ML) algorithms, despite the subtle biochemical differences between specimens with different ages (17). Both Near-Infrared Spectroscopy (NIRS, wavenumbers in the range 10,000 to 4000 cm^-1^) and Mid-Infrared Spectroscopy (MIRS, from 4,000 to 400 cm^-1^) have been proposed for age-grading Afrotropical malaria vector species of the *Anopheles gambiae* complex (18,19), as well as for species identification (19), pathogen detection in vectors (20) and humans (21) and blood meal identification (22). Both NIRS and MIRS have been also successfully tested for age-grading *Ae. aegypti* and for detection of *Wolbachia* infection in laboratory samples of this species (23,24). Overall, MIRS shows a higher mechanistic robustness as it provides distinct and well-defined absorption bands corresponding to fundamental molecular vibrations, crucial to enable the generalisation of age-grading predictions across different settings (17,25).

Less effort has been so far invested on the development of age-grading secondary vectors, such as *Ae. albopictus*, despite in the last decades, this highly invasive species has increased its public health relevance at the global level. In fact, in addition of being a secondary dengue virus vector and a primary one of Chikungunya virus in tropical regions, it has been responsible of the first autochthonous cases and outbreaks of these arboviruses in Europe (26–28). We could not find published research that measured the accuracy of morphological age-grading methods in this species, and attempts made in Sapienza University to test these approaches on laboratory-reared females of known age revealed very low accuracy (data not shown). Preliminary attempts on *Ae. albopictus* age-grading by NIRS (29) showed high accuracies with specimens raised in the laboratory, but not with field collected samples (30,31). To our knowledge, MIRS has never been tested for age-grading *Ae. albopictus*.

Here, we report the results of the first steps towards the development of a MIRS-ML framework that can be used to estimate the age of wild-caught *Ae. albopictus*. We demonstrated the ability of MIRS-ML to age male and female mosquitoes under laboratory as well as under natural conditions, and tested the capacity of the approach to detect age-structure shifts in *Ae. albopictus* populations in a simulated vector control intervention and to provide plausible age structures in field-collected males and females.

## MATERIALS AND METHODS

### Adult *Aedes albopictus* mosquito samples

We utilized three sets of *Ae. albopictus* adult samples in this study, i.e. laboratory-reared, semi-field reared and field-collected samples, as follows.

#### Laboratory-reared samples

Approximately 5,000 dried eggs from an *Ae. albopictus* colony maintained at Sapienza University were allowed to hatch and larvae were fed with fish food and yeast powder until pupation. Adults emerging in a single day were placed in a 30cm^3^ cage (300-600 adults/cage) and fed a 5% glucose solution. A total of 4 cages was obtained. Mosquitoes (from eggs to adults) were kept at 26 ± 1.0°C, 60 ± 5% RH, and a 14:10 photoperiod. Males and females were collected on day 1 post-emergence and subsequently every 7 days until day 15 and 36, for males and females, respectively (Table S1). To obtain variability in female gonotrophic stages, females in each cage were provided a blood-meal on different days before the collections. Defibrinated ram blood meals were provided by Hemotek Membrane Feeding System with 3-day intervals. Oviposition cups were supplied 2 days after each blood meal. This allowed us to collect females at various gonotrophic stages: unfed, freshly fed (1 day from bloodmeal), gravid (2 days), and post egg laying (3 days). Mosquitoes were killed using a chloroform-soaked cotton pad for at least 20 minutes to preserve the mosquito cuticle for infrared measurements (32). To avoid measurement alterations due to hydration differences, the mosquitoes were completely dried (17). For this purpose, specimens were placed in 15ml Falcon tubes with silica gel behind a cotton barrier and stored at ∼4°C until processing with MIRS.

#### Semi-Field Reared Samples

Mosquitoes were obtained from eggs collected via Ovitraps in Aprilia (LT, 41°35′40″ N, 12°39′15″ E) and the Rome metropolitan area (41°53′57.12″ N, 12°32′42.00″ E) in June and July 2023 to ensure a genetically heterogeneous population, following the rational implemented by Siria et al. (2022) (19). Rearing from hatching to adulthood and adult maintenance occurred at the Experimental Botanical Garden of Sapienza University (Rome, Italy). Approximately 10,000 field-collected eggs were divided into 32 basins (35×25cm), each containing 2 litres of water, maintaining a density of ∼1.5 larvae per ml. The larvae were fed daily with fish food and yeast powder until pupation. Larvae pupated over two consecutive days were separated into plastic cups (approximately 500 pupae/cup) and placed inside cages for emergence, so that each cage had mosquitoes emerging over a three-day period, for a total of 7 cages. Pupae not emerging in this time frame were removed. To obtain variability in female gonotrophic stages, females in each cage were provided a ram blood-meal at different days before collection via a Hemotek Membrane Feeding System brought directly into the semi-field site. Oviposition cups were supplied 2 days after each blood meal. Adults were collected every 3 days to obtain consecutive age classes (1-3, 4-6, 7-9, …, 31-33 days old) (Table S2). Average minimum, maximum and mean temperatures during the experiment were 28.5°C, 22.6°C, and 34.3°C, respectively. Mean relative humidity was 66.4% (33) and ∼14:10 photoperiod. Females were collected at various stages: unfed, freshly fed (1 day after bloodmeal, ABM), gravid (2 days ABM), and post egg laying (3 days ABM). The collection, killing, and storage procedures were consistent with those used for laboratory-reared mosquitoes.

#### Field-Collected Samples

Between May and June 2023, adult *Ae. albopictus* mosquitoes were sampled using BG traps placed in private gardens in Aprilia (LT, 41°35′40″ N, 12°39′15″ E). Mosquitoes were killed by exposure to chloroform for 20 minutes and stored in a similar way than the laboratory- and semi-field-reared specimens.

### Spectroscopy

Mosquito samples were measured using Bruker Alpha II FT-IR spectrophotometers equipped with a diamond ATR crystal at the Vector Biology Group laboratory (University of Glasgow, UK). Only the head and thorax were scanned by placing them at the centre of the crystal and pressing with the anvil attached to the instrument to maximize contact between the mosquito cuticle and the ATR crystal. Background and mid-infrared (MIR) spectra were acquired by averaging 24 scans at a resolution of 4cm^−1^ over a wavelength range of 4000–400cm^−1^. Low-quality spectra were discarded using a custom script in Python designed for mosquito spectra which consisted of three filters to: 1-eliminate spectra with atmospheric intrusion (CO_2_ and water vapour) by assessing the smoothness of the region between 3900 and 3500 cm^−1^; 2-eliminate low intensity spectra measuring the average absorbance of the plateau in the spectrum between 500 and 400 cm^−1^.; 3-eliminate distorted spectra caused by the anvil (17).

### Machine Learning analysis

Three datasets of infrared spectra were obtained: one from laboratory-reared individuals, one from semi-field samples, and one from wild populations. The analyses were conducted using Python 3.10 to develop a Supervised Machine Learning (ML) algorithm. The laboratory dataset was used to initially validate the ability of MIRS to detect spectral variations associated with different mosquito ages. The semi-field samples were then used to develop an ML algorithm capable of accurately estimating the age of *Ae. albopictus* individuals. This model was deployed at three different resolution levels, i.e. grouping spectra from mosquitoes emerged in 3, 6 or 9 consecutive days.

For lab and semifield datasets, we shuffled and split the dataset into the training (80%) and test sets (20%), stratified by age groups. The training set was standardised and used to compute baseline performance of eight ML algorithms – LR: Logistic Regression; RF: Random Forest; SVC: Support Vector Classifier; KNN: K-Nearest Neighbours; DT: Decision Tree; GB: Gradient Boosting Classifier; AB: Ada Boost Classifier; ET: Extra Tree Classifier. In all cases, 10-fold cross validation and the default parameter settings of the models were used. The best model was chosen based on accuracy as well as its suitability for analysis of high dimensional data. It was then optimised using hyperparameter tuning, which consists of choosing a set of optimal values for the model hyperparameters to maximise its performance. Hyperparameter optimisation was conducted using gridsearch with 10-fold cross-validation to identify the best-performing parameter combinations. The selected parameters for our optimised models are reported in Table S3 and Table S4. The remaining 20% of the data (the test set) were used for the final evaluation of the optimised models. Model performance was measured by comparing the estimated age with the known age of the specimens, resulting in a confusion matrix that reported accuracy for each age class. The final optimised model trained in semifield data was used to predict different age classes using data from wild mosquitoes. The analysis was performed using scikit-learn 1.2.2.

### Population simulation and power analysis

To estimate the statistical power of the MIRS-ML model to detect a shift in the age structure of a mosquito population after an insecticidal intervention, we generated computer-simulated mosquito populations under two different scenarios: (i) no intervention and (ii) an adult spray intervention with immediate action of 50% killing efficacy; populations were supposed to be sampled one week after the intervention. The age structure (i.e., the frequency of each age class) in the populations was simulated assuming a constant daily mortality of 4% up to 33 days for females and up to 15 days for males, with no survival after 33- or 15-days post-emergence, respectively. The age structure of each post-intervention population was compared with the control population using Wilcoxon/Mann–Whitney U tests for both females and males (high-resolution algorithm). For males analysed with mid- and low-resolution algorithms, a Fisher exact test was used instead, due to the smaller number of available age classes. Power was estimated as the proportion of 10,000 simulated datasets where a significant (p<0.05) difference in age structure was detected between intervention and control populations across seven sample sizes (n = 20, 50, 100, 150, 200, 250 and 300). Simulations were performed in R v4.3.2.

## RESULTS AND DISCUSSION

### Mid-infrared spectra of laboratory-reared *Ae. albopictus* varies across ages

To test whether MIRS could be used to age-grade *Ae. albopictus* populations, we first applied it to a laboratory dataset. Specifically, we reared and collected 703 female and 316 male adults at different ages (1-, 8-, 15-day old for both females and males, and additionally 22-, 29- and 36-day old for females only) and measured them with MIRS (Table S1). We then compared eight different algorithms in terms of their ability to estimate the age of these samples based on their MIR spectra. The models ranked similarly for males and females independently. In both cases, Support Vector Classifier (SVC) had the highest baseline accuracy followed closely by Logistic Regression (LR), which has been used before in similar applications (17). The baseline accuracies of SVC were 76.1% and 94.8% for females and males, respectively (Figure 1A, C).

**Figure 1:**
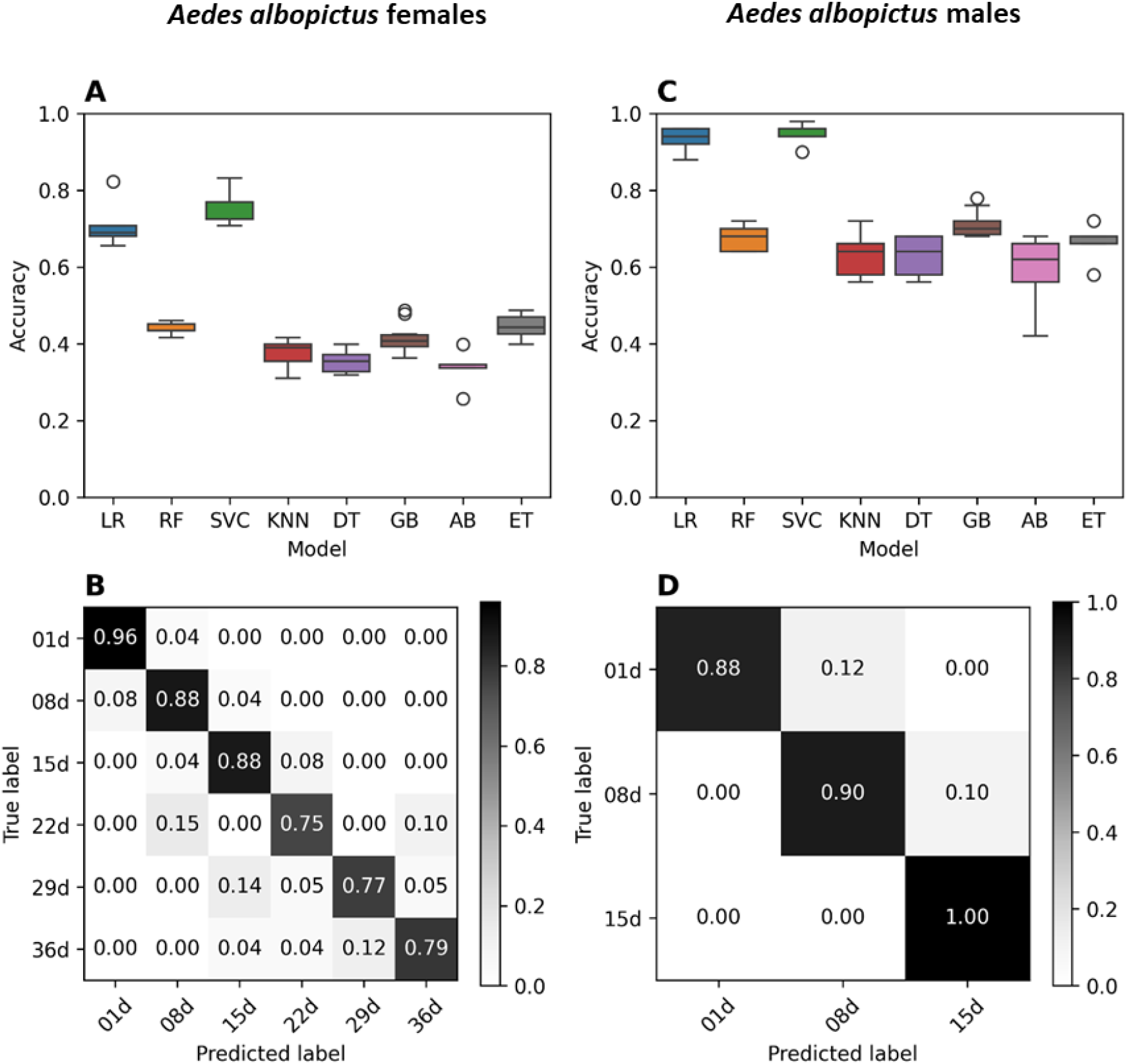
MIRS-SML age prediction for *Ae. albopictus* adults reared in laboratory. **A, C**: Boxplots showing accuracy for each tested model for females (**A**) and males (**C**). The colour bar on the left shows the scale of colours that correspond to each accuracy value: from white (0, lowest accuracy) to black (1, highest accuracy). LR: Logistic Regression; RF: Random Forest; SVC: Support Vector Classifier; KNN: K-Neighbours; DT: Decision Tree; GB: Gradient Boosting Classifier; AB: Ada Boost Classifier; ET: Extra Tree Classifier. **B, D:** Confusion matrices of age classification for adult females (**B**) and males (**D**). Y axis shows the known ages of mosquitoes, while x axis shows the ages predicted by SVC model. Each cell of the matrix shows the percentage of individuals assigned to each age class. Percentages on the diagonal correspond to individuals whose age was correctly.

As initially the algorithms trained using the whole mid-infrared spectrum (4,000–400 cm^−1^) tended to learn from chemically irrelevant regions (mainly 2,500–1,800cm^−1^, Figure S1), we preselected the region between 1800 and 402 cm^−1^ for the final model. This pre-processing step, also present in similar applications like in Pazmiño Betancourth et al. (2023) (34), is designed to reduce the noise-to-signal ratio of the spectral data. By restricting the data available to the models to only those regions of the spectra with meaningful chemical information, we reduce the likelihood of overfitting and increase the generalisability of our models.

Prediction accuracy of each age class ranged from 75% to 96% for females (up to 36 days old) and from 88% to 100% for males (up to 15 days old, Figure 1B, D). The confusion matrix in Figure 1B also illustrates that, when incorrect, the model generally predicted ages close to the true value. Together, these results demonstrate that the MIR spectra carry a signal that correlates with the age of the individual, and that this signal could be detected by the model, supporting the potential of the MIRS-ML approach as an age-grading method for wild *Ae. albopictus*.

### Models based on MIR spectra from genetically and environmentally variable *Ae. albopictus* can accurately predict age classes

To develop a MIRS-ML approach capable of predicting the age of mosquito field populations, it is essential to train models on spectra from mosquitoes reared outdoors under natural conditions and representing real genetic backgrounds, ensuring that both environmental and genetic variability influencing ageing are properly captured. To this aim, approximately 10,000 field-collected *Ae. albopictus* eggs were reared under semi-field natural conditions; adults were collected at different ages and gonotrophic stages. The final dataset comprised a total of 1,225 and 565 MIR spectra for females and males, respectively (Table S2).

For each sex we developed three machine learning models at high (i.e. successive age classes, each of which including adults emerged over a 3-day period), medium (age classes including adults emerged within 6 days), and low resolution (age classes including adults emerged within 9 days) (Table S5). Similarly to spectra from laboratory reared mosquitoes, SVC had the highest accuracy (Figure S2) and was used for further optimisation.

The prediction accuracy decreased as the resolution increased. The highest overall model accuracy was obtained with the low-resolution trained model (89.4% for females and 99% for males), then medium-resolution (78.5% for females and 93% for males), and finally the high-resolution (72.6% and 85.8%, respectively) (Figure 2). Generally, the youngest age class had higher prediction accuracies, suggesting that changes in cuticle composition are larger during the first days after emergence. This was also observed on *Anopheles* malaria vectors (19). The lower accuracy in predicting female age compared to males may be due to greater physiological variability, as the dataset included unfed, blood-fed and gravid females. Notably, in all cases, but particularly in the high-resolution model, when age classes were misclassified, the model tended to predict the nearest age classes, and not just random ages.

**Figure 2.**
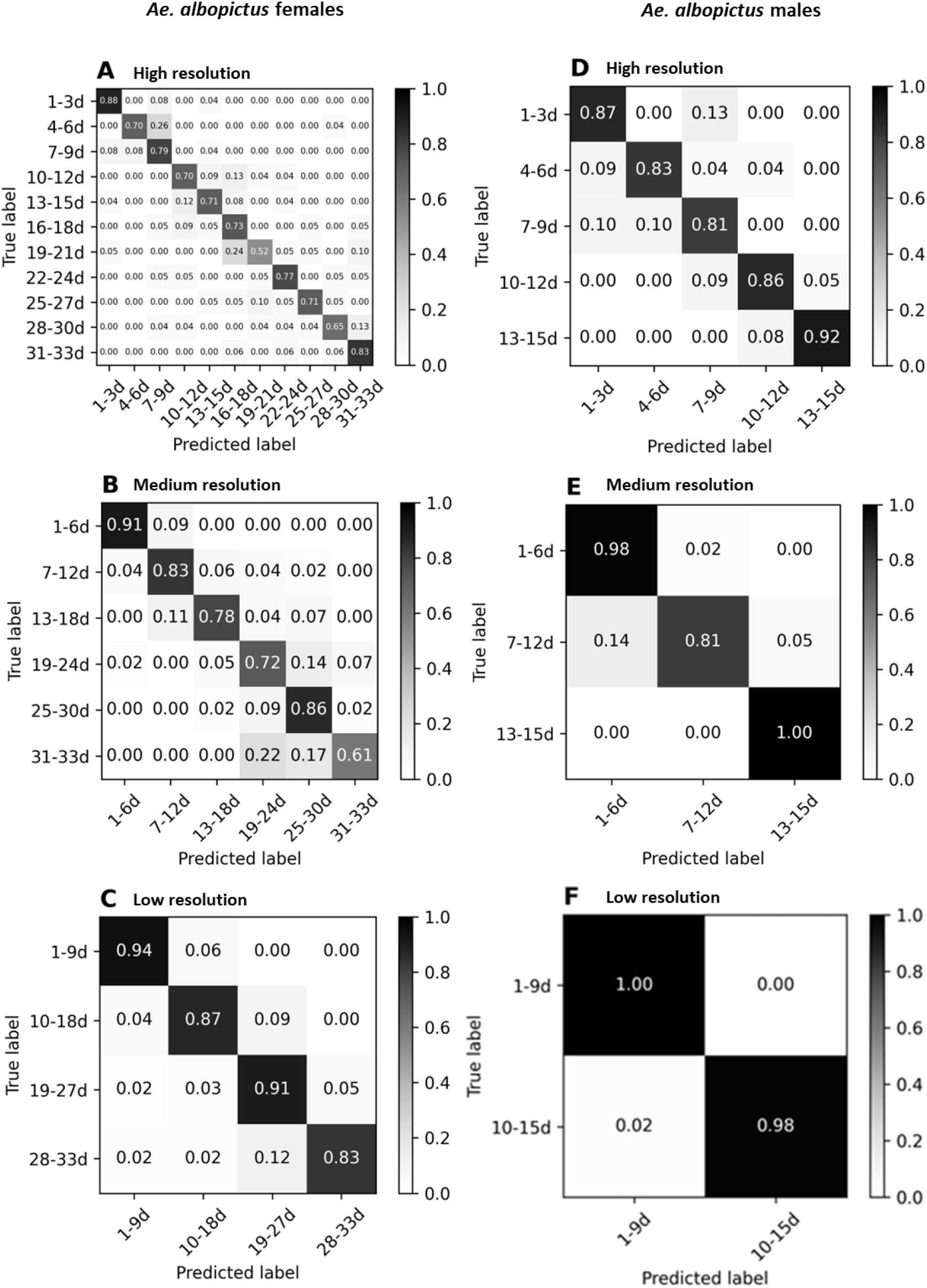
Confusion matrices for three different age classification resolutions for *Ae. albopictus* females (left) and males (right), reared under semi-field conditions. **A**,**D**: high resolution (3-day classes); **B**,**E**: medium resolution (6-day resolution); **C**,**F**: low resolution (9-day classes). Y axes show the true ages of mosquitoes, while X axes show the ages predicted by SVC algorithm. Each cell of the matrix shows the accuracy (between 0 and 1) of individuals assigned to each age class. Percentages on the diagonal correspond to individuals whose age was correctly estimated by the algorithm. The colour bar on the left shows the scale of colours that correspond to each accuracy value: from white (0, lowest accuracy) to black (1, highest accuracy).

### MIRS machine learning models learned from sex-specific chemical features

To better understand which spectral regions contributed most to the classification, we generated feature importance plots based on the model coefficients (Figure 3, Table S6, Table S7). The plots display the mean mid-infrared spectrum alongside vertical lines indicating the top 50 wavenumbers with the highest mean absolute coefficient values. These highlighted features represent the spectral regions from which the model learned the most to discriminate between age classes. By comparing the feature importance plots within the same sex, some regions seemed to be consistently influential (Figure 3).

**Figure 3.**
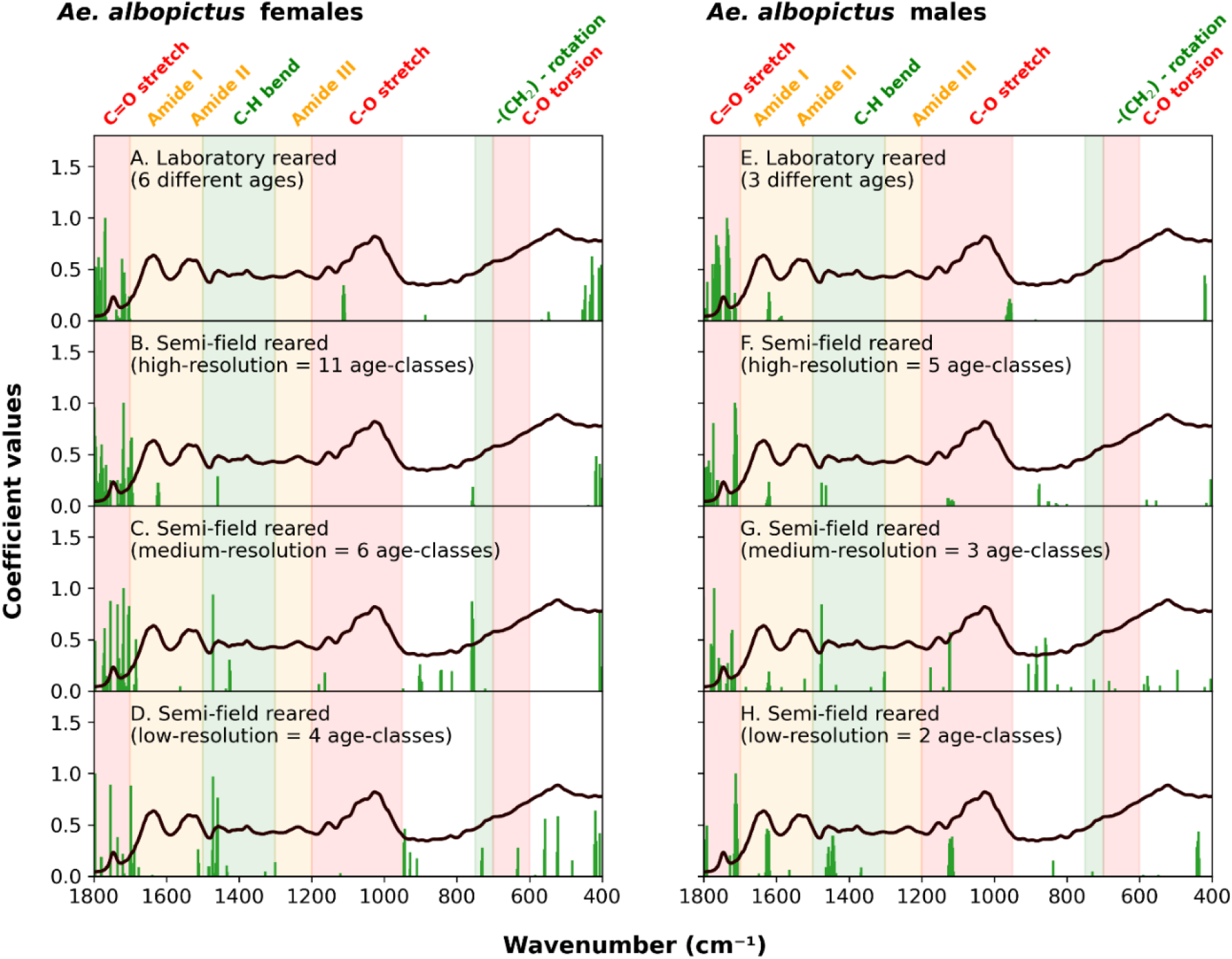
Mean mid-infrared spectrum in the 400-1800 cm^-1^ range of *Aedes albopictus* adult. mosquitoes, overlaid with the top 50 most important wavenumbers (features) identified by the SML model. Vertical green lines indicate the wavenumbers with the highest absolute model coefficients values. These features represent the most influential bonds in predicting mosquito age. (**A**) females reared in laboratory; (**B**) semi-field females at high resolution; (**C**) semi-field females at medium resolution; (**D**) semi-field females at low resolution; (**E**) males reared in laboratory; (**F**) semi-field males at high resolution; (**G**) semi-field males at medium resolution; (**H**) semi-field females at low resolution.

Features located at 1746 cm^-1^ (corresponding to C=O stretches related to wax, protein and chitin) were the most influential for age classification regardless of sex and rearing conditions (Figure 3A-H). Similar features have been observed in MIRS-ML models that predict the age of other vectors, specifically in the female malaria mosquitoes *An. gambiae* s.l. (19) and *An. funestus* (35) and in the female tsetse flies (36).

In addition, other wavelengths became influential as the age resolution of semi-field reared samples decreases. In females, these are mostly C-H bend (1457 cm^-1^ and ∼758 cm^-1^) related to chitin, wax and proteins (Figure 3B-D). In males (Figure 3F-H), other biological relevant areas are amide I (1636 cm^-1^, related to chitin and protein) C-H bend (1457 cm^-1^ related to wax and proteins) and C-O stretch (1100 cm^-1^, related to chitin, waxes and proteins).

### MIRS-ML can detect *Ae. albopictus* age structure shift following a simulated intervention

Age-grading vector populations allows the detection of shifts in mosquito age structures after an intervention, providing a direct way to quantify its impact. We tested if our semi-field MIRS-ML predictions were sufficiently accurate to detect an age-structure shift and compared the models at different age resolutions. First, we simulated the age structure of 1) a control population (4% daily mortality) and 2) a population sampled one week after an adulticide intervention of 50% efficacy. Then, we applied our MIRS-ML models to reconstruct the predicted proportions of different age classes and compared with 100% correct predictions (Figure 4A-B, Figure S3A-E). Finally, we estimated the statistical power of MIRS-ML to detect a shift in the different age classes expected from the intervention (Figure 4C, Figure S3C-F). When applying MIRS-ML on females using the high-resolution age classes, sampling less than 100 mosquitoes pre- and post-intervention was sufficient to obtain >80% power to detect an age structure shift; however, when using medium- and low-resolution classes a larger sample size was required to achieve >80% power, showing that predicting more age classes, even if with overall lower accuracy, is the most powerful approach to detect a population age structure shift following a vector control intervention. Similar results were obtained when the models were applied to male mosquito populations (Figure S4).

**Figure 4.**
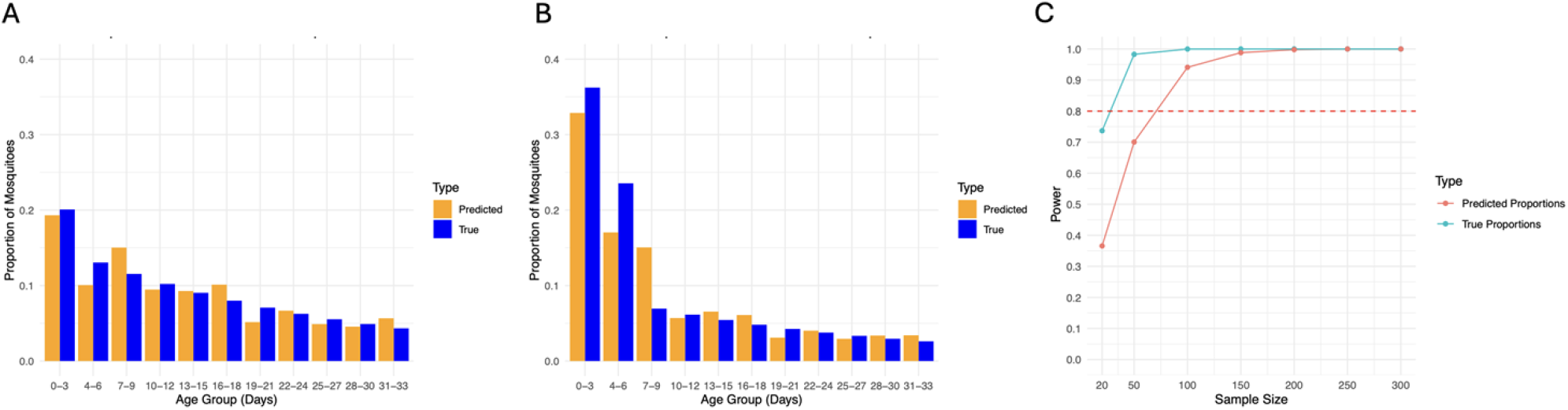
Modelling the assessment of the effectiveness of vector control intervention by shifts in population age structures detected by MIRS-SML. Computer simulations were used to assess the power of MIRS-SML model to detect an age structure shift between **A)** an *Ae. albopictus* natural population with 4% daily mortality relative to **B)** a population target of a control intervention killing 50% of the females 1 week earlier. Blue and orange bars indicate the simulated age structure and predicted age structure based on the MIRS-female-high-resolution model, respectively. **C)** Power to detect an effect of the vector control intervention was estimated over seven sample sizes per population from 20 to 300. The blue line shows the power that would be achieved with 100% accurate age group classification and the red line indicated the power using the MIRS model. The dotted line indicated 80% power at p<0.05.

### MIRS-ML predicted plausible age structures from both laboratory-reared and wild collected *Aedes albopictus* female and male adults

We then evaluated the generalisability of the low-resolution semi-field MIRS-ML model by testing it on two independent, previously unseen datasets. This was not carried out with the higher resolution modes due to their sparse age distribution.

First, the model was applied to the laboratory dataset. Notably, the model correctly assigned most of 1 and 8 day old females to the first age class (1-9d), overestimated the assignment of 15 and 22 old females to the intermediate age classes, and largely underestimates the assignment of 29d old females to the oldest age class (Figure 5A). In the case of males – for which only 2 age classes were considered – the model underestimated the younger class (Figure 5B). One possible explanation for the underestimation of age in laboratory-reared females is that the semi-field MIRS-ML model was trained on females exposed to more natural environmental conditions including warmer temperatures and larger daily thermal excursion, which likely accelerated their biological ageing. As a result, older laboratory-reared females may be biologically younger than their semi-field counterparts of the same chronological age. A similar pattern was reported in *Anopheles gambiae s*.*l*. females, where models trained on laboratory data overestimated the age of semi-field mosquitoes (19). The reason for the overestimation of age in laboratory-reared males is less clear. One possibility is that males aged more rapidly in the laboratory due to more frequent mating opportunities, which are typical in colony conditions.

**Figure 5.**
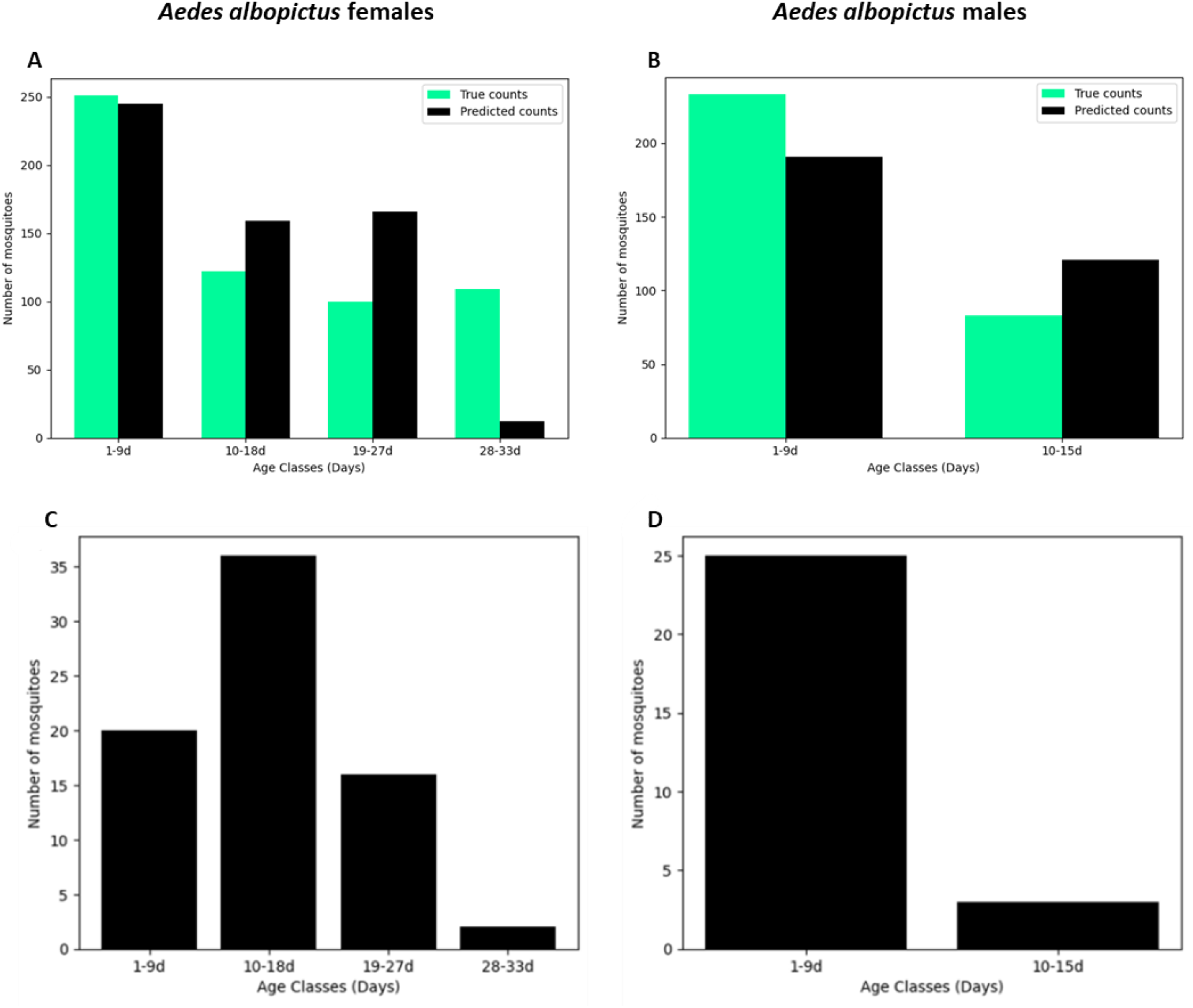
MIRS-ML age structures predictions of lab and wild collected *Aedes albopictus* adults. Barplots show the number of *Aedes albopictus* lab females (**A**) and males (**B**) and wild females (**C**) and males (**D**) assigned to different age classes bases on MIRS-ML low resolution model. **A** and **B**: laboratory-reared females were 1, 8, 15, 22 and 18 days old; laboratory-reared males were 1, 8 and 15 days old. **C** and **D**: females and males of unknown ages.

Then, we tested the model on 74 female and 28 *Ae. albopictus* males sampled in the field near Rome, which, having experienced natural climatic conditions, could be expected to exhibit more comparable ageing rates. As determining the number of gonotrophic cycles by examining the ovariole dilatations is not feasible in this species, our test simply aimed at assessing if the predicted age structure would be realistic in field populations. Our predictions reconstructed age structures that are plausible in relation to the expected ones (Figure 5C, D). In females, the majority (48%) was classified as medium old (10-18 day old), the rest as either younger or older (49% combining 1-9 days and 19-27 days) and a very small proportion (3%) as the oldest age class (28-33 days). In males, the large majority (89%) was classified as the youngest age class (1-9 day old). These results also have implications for disease transmission. Indeed, assuming that in the presence of arbovirus circulation females >9 day old would have had time to feed on an infected host and to allow completion of the virus extrinsic incubation, this would imply that >75% of the local population had the potential to transmit the virus.

## CONCLUSION

The results obtained represent a first relevant step towards the development of a sound and reproducible MIRS-ML approach for age-grading of *Ae. albopictus* populations in their invasive range. First, we demonstrated the ability of MIRS-ML to age-grade male and female mosquitoes under laboratory conditions. Second, we optimised the model with mosquitoes reared under natural conditions in a semi-field facility, in order to expose them to more realistic ambient conditions. Then, we showed that the models are capable of detecting age-structure shifts in *Ae. albopictus* populations in a simulated vector control intervention. Finally, we validated MIRS-ML on unseen data and reconstructed plausible age structures in field-collected *Ae. albopictus* male and female mosquitoes.

Future work will focus on improving the method’s accuracy and generalisability by expanding the training datasets with spectra from more diverse mosquito populations and incorporating environmental variables known to influence mosquito ageing. While our developed model demonstrated some generalisability when applied to unseen datasets—including both laboratory-reared and field-caught *Ae. albopictus* mosquitoes - it has not yet been tested on more genetically and environmentally diverse samples. Since MIRS detects age-related changes in the mosquito cuticle (17), variation in genetic background or environmental conditions may affect the resulting spectra and, consequently, the performance of MIRS-ML models. In our study, the field validation—though it produced plausible age structures—was based on a population genetically similar to the one used for training. Moreover, both the training and field-sampled mosquitoes were collected from the same region and during the same season, likely experiencing similar environmental conditions. Therefore, while our results are promising, the current MIRS-ML models require broader validation and refinement across diverse populations and ecological contexts to become a robust tool for global vector surveillance.

Due to the low processing costs, we expect this approach to have the potential to be largely exploited in the future to characterise the age structure of *Ae. albopictus* field populations for improved epidemiological risk models and for assessment of the effectiveness of mosquito control interventions. This is particularly valuable in non-endemic settings where *Aedes albopictus* has become established, as the impact of interventions cannot be directly measured through changes in disease prevalence. By focusing on the age structure of the population in an area treated with vector-control, it would be possible to gain a deeper understanding of how adulticide interventions affect mosquito populations over time and their contribution to target the older, disease-transmitting vectors, providing a more comprehensive assessment of their effectiveness (37,38).

## AUTHOR CONTRIBUTIONS

Conceptualization: AdT, BC, FB

Mosquito rearing: MF, MM, PS

MIRS-spectra measurements: MF, MPB, IC

Data analysis and modelling: MF, MPB, FB, BC

Writing – original draft: AdT, FB, MF

Writing – review & editing: all authors

## DATA

Data can be accessed at: https://github.com/casasgomezuribarri/Foti_et_al_albopictus_DATA Scripts can be accessed at: https://github.com/maurocolapso/AlbopictusMIRS_Foti_et_al_2025.git

